# Inferring spatially transient gene expression pattern from spatial transcriptomic studies

**DOI:** 10.1101/2020.10.20.346544

**Authors:** Jan Kueckelhaus, Jasmin von Ehr, Vidhya M. Ravi, Paulina Will, Kevin Joseph, Juergen Beck, Ulrich G. Hofmann, Daniel Delev, Oliver Schnell, Dieter Henrik Heiland

**Author notes:** Equal contributed first authorship (Jan Kueckelhaus, Jasmin von Ehr). **Corresponding author:** Dieter Henrik Heiland, Department of Neurosurgery, Medical Center University of Freiburg, Breisacher Straße 64, 79106 Freiburg, -Germany.

## Abstract

Spatial transcriptomic is a technology to provide deep transcriptomic profiling by preserving the spatial organization. Here, we present a framework for SPAtial Transcriptomic Analysis (SPATA, https://themilolab.github.io/SPATA), to provide a comprehensive characterization of spatially resolved gene expression, regional adaptation of transcriptional programs and transient dynamics along spatial trajectories.

## Brief Communication

Deep transcriptional profiling of single cells by RNA-sequencing maps the cellular composition of tissue specimens regarding cellular origin, developmental trajectories and transcriptional programs^1–3^. However, information determining the spatial arrangement of specific cell types or transcriptional programs are lacking and thus can only be predicted indirectly^4^, which is a considerable drawback of this method. Spatial tissue organization was traditionally investigated by imaging technologies which provide information at high resolution but are strongly limited by the number of genes or proteins to be mapped. Several novel technologies such as MERFISH^5^, FISH-seq^6^, Slide-seq^7^ or spatial transcriptomics^8,9^ are able to preserve the spatial context of transcriptional data, however all these technologies are limited by either the spatial resolution or depth of transcriptional profiling. Further, data integration, visualization and analysis of transcriptomic and spatial information remains challenging. Here, we present a software tool to provide a framework for integration of high-dimensional transcriptional data within a spatial context. By combining user-friendly interfaces for visualization, segmentation or trajectory analysis and command-based pipe-friendly functions for data manipulation and modeling, we provide a broad range of applications for different analytical demands. In addition, we implemented interfaces to provide easy exchange of numerous external tools. Previously published tools focus mainly on the visualization of gene expression using known tools from scRNA-seq analysis rather than addressing gene expression within its spatial context^10–12^. In particular, we focus on transient changes of gene expression and aim to infer transcriptional programs that are dynamically regulated as a function of spatial organization.

In order to present an overview of possible analytic capabilities of the SPATA workflow, Figure 1a, we generated spatial transcriptomic datasets from human cortex and human glioblastoma samples using the Visium technology (10X Genomics). The human cortex is separated into defined layers containing different types of neurons and cellular architecture. In a first step, we combine shared-nearest neighbor clustering and spatial pattern recognition by an external tool (spatial pattern recognition via kernels, “SPARK”^13^) in order to determine genes with a defined spatially resolved expression pattern. We found that the cortical layering is accurately reflected by our clustering approach. In order to gain insights into the spatial organization we provided a tool to compute the spatial distance within the defined layers or correspondent clusters. An increasing distance within individual clusters allows to differentiate between narrowly related or a widespread dispersion of spots within the cluster.

**Figure 1:**
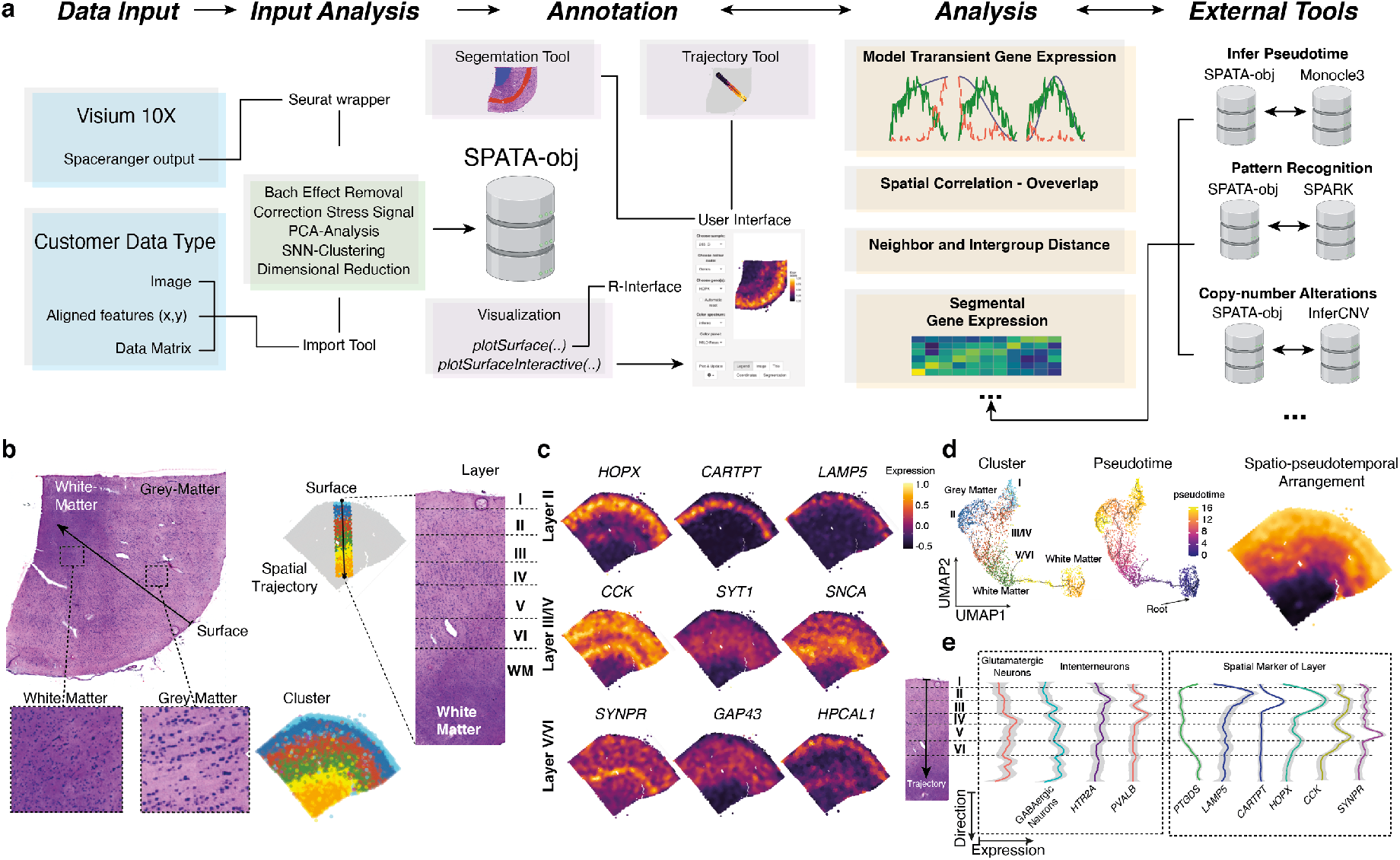
a) Illustration of the SPAtial Transcriptomic Analysis (SPATA) workflow containing Data Input and a predefined set of analysis which will be saved in a SPATA object. In the following step of the workflow, annotation of segment trajectories can be performed using a user-friendly interface. Additionally, multiple tools for visualization are available within the user interface. This also includes tools for geneset enrichment analysis (GSEA) and gene set variation (GSVA). After region of interests are defined, a list of analytic tools is provided which includes wrapper for external tools. b) H&E staining of a human cortex sample with the corresponding SNN clustering (bottom right). The annotation tool is used to draw a trajectory along all cortical layer (right side) c) Infer genes with defined peaks along the trajectory revealed genes with layer specific gene expression. d)Integrating an external package (monocle3) the pseudotime within the sample was computed and visualized by a 2D UMAP representation (left) and within its spatial context (right). e) Comparison of traditional markers (left) and markers given by our model (inferring transient gene expression) along the cortical layering.

Next, the spatial overlap of transcriptional programs or gene expression was analyzed using a Bayesian approach, resulting in an estimated correlation which quantifies the identical arrangement of expression in space. In a further step, we aimed to analyze dynamic changes, which were annotated using pseudotime estimation or RNA-velocity. We directly implemented the pseudotime inference from the monocle3^14^ package, but also allow the integration of any other tool such as “latent-time” extracted from RNA-velocity (scVelo^15^). Another option for dynamic gene expression analysis is the detection of defined transcriptional programs along a defined trajectory. In our example, we mapped different activation states of astrocytes and microglia within the cortical layering.

Moreover, we provide the opportunity to screen for gene expression or transcriptional programs which transiently change along predefined trajectories by modelling gene expression changes in accordance to various biologically relevant behaviors. All genes or transcriptional programs which significantly followed one or multiple predefined models were ranked and visualized. The detection of dynamic spatially defined gene expression patterns is also of great interest in malignant specimens. In another example, we profiled tissue of a human glioblastoma, the most malignant tumor of the central nervous system (CNS) as SPATA provides numerous tools to analyze datasets with malignant origin. In a first step, integrating inferred copy-number alterations (CNV)^2,16^, spatial pattern recognition and shared-nearest neighbor clustering provides a broad overview of spatially defined transcriptional programs within the subclonal architecture of tumor samples, *Figure 2a-g*. Using this information, specific segments can be specified and analyzed to gain insights into their spatially differentially expressed genes. We showed that segments of higher cellular density also contained increased signaling of the hypoxic pathway including expression of *VEGFA*, *HIF1A* and *GAPDH*. Additionally, mapping the subclonal architecture based on a CNV clustering allowed to screen for gene expression differences within regions of exclusive genetic context *Figure 2d*. Inferring spatially transient gene expression along trajectories connecting particular tumor regions, i.e. between tumor core and infiltration zone, provided the opportunity to map transcriptional programs executed during tumor infiltration and tumor-induced microenvironment changes of the surrounding areas. Thus, we were able to show that immune related genes from myeloid cells and reactive astrocytes were localized in a “glial-scars” resembling structure, sharply separating normal brain from tumor regions *Figure 2h*. We observed a transient increase of macrophage and microglia activation directed towards the tumor boarder. Mapping transcripts that mark for lymphoid cells, we found more T cells abundance within the normal brain compared to tumor regions which is in line with the reported immunosuppressive environment within glioblastoma. Inferring pseudotime, we were able to confirm a dynamic adaptation of myeloid cells along our defined trajectory *Figure 2i-j*. Recently, Neftel and colleagues established a classification of 4 transcriptional states using single-cell RNA-sequencing, *Figure 2f,i*. Using these signatures, we were able to map the spatial distribution of assumed tumor heterogeneity. We implemented a 2D representation of all 4 states which could be used to map the distribution of all transcriptional states within defined segments or along spatial trajectories. Of utmost importance, our tool enables the usage of a variety of different biological data containing a spatial context such as spatially resolved mass-spectroscopy or imaging mass cytometry (IMC). SPATA is a resource developed from scientists for scientists incorporating the FAIR principles of providing findable, accessible, interoperable, and reusable data^17^.

**Figure 2:**
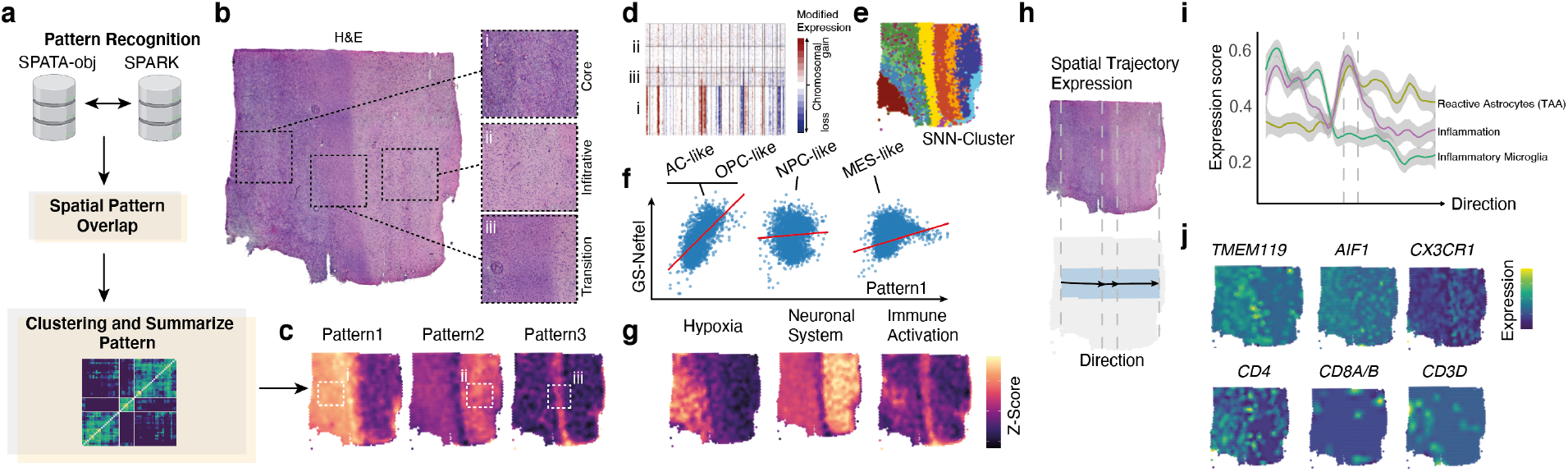
a) Illustration of the SPATA workflow integrating SPARK for pattern recognition into the analysis of human glioblastoma. SPARK will estimate to what extent a gene is present in a spatial pattern. The output is piped into a spatial overlap analysis and clustered to extract set of genes which belong to the same pattern. b) H&E staining of glioblastoma with 3 histological distinct regions. c) Predicted pattern visualized by the z-scored gene expression of all genes aligned into a pattern. d-e) Copy-number analysis of the sample and cluster annotation (e). f) Comparison of recognized pattern with known gene expression classification, here the Neftel classification. g) Expression of significantly expressed pathways within a pattern. h) Spatial trajectory analysis along the tumor infiltration region. i-j) Change of z-scored geneset expression along the trajectory (i) and marker genes of microenvironmental alterations and inflammation (j).

## Methods

### SPATA software and functions

A detailed overview of all included functions and the structure of the package is given at the package website (https://themilolab.github.io/SPATA/index.html). We implemented tutorials for all described analytic approaches to provide a simple-as-possible solution to trace the individual analytic steps.

### Data preparation, per-analysis and SPATA object implementation

We offer two possible input options. On one side, we implemented the direct input from spaceranger by using the Seurat wrapper for spatial transcriptomics. On the other hand, we used the Seurat v3.0 package to normalize gene expression values by dividing each estimated cell by the total number of transcripts and multiplied by 10,000, followed by natural-log transformation. As described for single cell-RNA sequencing, we removed batch effects and scaled data using a regression model including sample batch and percentage of ribosomal and mitochondrial gene expression. For further analysis we used the 2000 most variable expressed genes and decomposed eigenvalue frequencies of the first 100 principal components and determined the number of non-trivial components by comparison to randomized expression values. The obtained non-trivial components were used for SNN clustering followed by dimensional reduction using the UMAP and TSNE algorithm. After analysis all date will be saved in a SPATA object, detailed information of the S4 object structure is given at the package information. Another option is to provide 3 files that will be used to create a SPATA object, one file containing barcode information or other identifier of each spot with the given x and y coordinates determining the spatial position of each spot within the H&E image. The second file contains an expression or intensity matrix with identifier as colnames and genes or other features as rownames. The last file is an image with x and y coordinates corresponding to the identifier of file1. If the inputs are gene expression counts we run the standard pipeline (Seurat wrapper), otherwise (IMC, MALDI or MERFISH) we provide a data analysis pipeline which is designed for non-integer inputs and normal distributed data.

### Modeling of transient gene expression along spatial trajectories

A given trajectory includes multiple spots summarized into predefine bins of the directed trajectory. In order to model the gene expression of single genes or genesets we created a set of mathematical models which represent defined biological behaviors, including linear, logarithmic or gradient ascending/descending expression pattern, one-, or multiple peak expression, detailed information in the package description. The analysis is implemented into the function `assessTrajectoryTrends()`. Further, if a defined pattern is requested, we open the possibility to add a vector containing the requested model for which the algorithm will screen. Next, we fitted the summarized expression values of each bin using a non-parametric kernel estimation (Gaussian or Cauchy-Kernel), input vectors were normalized and z-scored: (1) 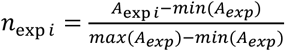 (2) 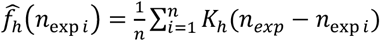 K is the kernel and 0.7 > h > 0.3 is used to adjust the estimator. Next, we computed residuals for each input vector (gene expression) and estimated area under the curve (AUC) using the trapezoidal numerical integration. (3) 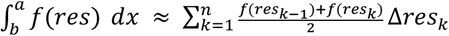 The distance and direction is defined by [a,b] a=x_0_ < x_1_<,…< x_n−1_ <x_n_=b. We use the AUC to rank the estimated models and predict genes that follow our predefined behavior. The implemented function `plotTrajectoryFit()` shows the model fit with respect to the given residuals.

### Enrichment analysis for SPATA

Gene sets were obtained from the database MSigDB v7 and internally created gene sets are available at within the package. For enrichment analysis we provide multiple methods listed in the description of the `plotSurface()` function. Per default, we use a probability distribution fitting of the input values which could be genes or summarized gene sets and transformed the distribution to representative colors. Further adaptation of the applied color scale can be performed by using the `confuns::scale_color_add_on()`. Further additions for geneset enrichment analysis or gene set variation analysis are implemented by using the GSVA package. As input for a GSEA the normalized and centered expression data are used and further transformed to z-scores ranging from 1 to 0. Genes were ranked in accordance to the obtained differential expression values and used as the input for GSEA.

### Two-dimensional representation of cellular states

Within the SPATA toolbox, we allow to plot a recently popular 2D presentation of multiple cellular states. As usual in SPATA, we provide two versions to acquire the data, on one side plotting from inside a SPATA-object is possible (`plotFourStates()`) and on the other hand, data can be used from outside (for example an expression matrix containing a single-cell dataset) by applying version 2 of the function(`plotFourStates2()`). Therefore, we aligned spots to variable states based on defined gene sets: GS_(1,2,..n)_. We separated cells into GS_(1+2)_ versus GS_(2+4)_, using the following equation: *A*_1_ = ‖ *GS*_(1)_, *GS*_(2)_ ‖_∞_ − ‖ *GS*_(3)_, *GS*_(4)_ ‖_∞_ A1 defines the y-axis of the two-dimensional representation. In a next step, we calculated the x-axis separately for spots A1<0 and A1>0: 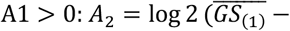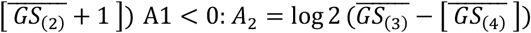 For further visualization of the enrichment of subsets of cells according to gene set enrichment across the two-dimensional representation, using a probability distribution fitting - we transformed the distribution to representative colors. This representation is an adapted method published by Neftel and colleges recently^2,3^.

### Spatial distance measurement

In order to measure the spatial distance, we use either a defined factorized input or a continuous vector. We fist measure the spatial distance from each spot to all other spots and compute a distance matrix with spots as rows and columns (n_r_=n_c_). If factorized input was applied, we factorize the matrix and calculate the mean distance per factor (f_1−>_ f_i_): (1) 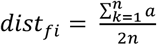 (2) 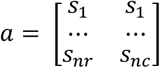. If a distance is numeric, we created bins of spots with common gene expression of gene set enrichment resulting in factorized values. Using the distance computation, we estimate to what extent a gene is expressed in exclusive spots (lower distance) or diffuse without spatial enrichment.

### Spatial overlap and correlation analysis

Spatial overlap of spatial correlation was designed to estimate the similarity of gene expression pattern within the spatial organization. In order to map spatial correlated gene expression or gene set enrichments, we used z-scored ranked normalized expression values. We used a Bayesian approach to compute the correlation distribution within two different genes or gene sets (~5-20 minutes runtime, MacOS 2019). The spatial reference is given by the x and y coordinates of each spot. In order to provide an alternative method which is computationally less-intensive (~1-3 minutes runtime, MacOS 2019) we construct a trajectory of spots from lowest ranked to highest ranked spot (based on z-scored input vectors). The genes of interest (which were correlated with the spatial trajectory) were fitted by loess-fit from the stats-package (R-software) and aligned to the ranked spots and fitted by a probability distribution. Correlation analysis was performed by Pearson’s product moment correlation coefficient. For heatmap illustration the gene order was computed by ordering the maximal peak of the loess fitted expression along the predefined spatial trajectory.

### Implantation of external tools: SPARK

For pattern recognition of spatially distinct expressed genes we integrated the R package SPARK^13^, which was shown to perform beneficial compared to other tools such as SpatialDE^18^. We transformed the required data into a SPARK object which is externally analyzed and reimported to SPATA. We add the possibility to group genes with a significant spatial pattern by overlap estimation and SNN clustering of the given correlation matrix.

### Implantation of external tools: InferCNV

Copy-number Variations (CNVs) were estimated by aligning genes to their chromosomal location and applying a moving average to the relative expression values, with a sliding window of 100 genes within each chromosome, as described recently^16^. First, we arranged genes in accordance to their respective genomic localization using the CONICSmat package (R-software). As a reference set of non-malignant spots, we used cortex from epilepsy patients. To avoid the considerable impact of any particular gene on the moving average we limited the relative expression values [−2.6,2.6] by replacing all values above/below *exp_(i)_*=|2.6|, by using the infercnv package (R-software). This was performed only in the context of CNV estimation as previously reported^19^.

### Implantation of external tools: Monocle3 or RNA-velocity

We implemented a wrapper to easily switch between cds-objects (monocle3) and SPATA objects. First, we compute minimum spanning tree (MST) to estimate the most separate paths and order these cells to annotate pseudotime. By using the `createPseudotime()` function, a shiny-interface from monocle3 will give the possibility to select a root for pseudotime annotation. Further, we provide the possibility to implement each vector, for example “latent time” extracted from RNA-velocity using scvelo, to integrate into our SPATA object.

### Data acquisition of spatial transcriptomics

All Visium Gene Expression experiments were performed according to 10X Genomics user guide ‘Visium Spatial Gene Expression Reagent Kits’. In brief, 10μm thick, cryosectioned slices of fresh frozen brain tissue were applied onto capture areas of Visium Spatial Gene Expression Slide, hematoxylin and eosin stained and imaged for subsequent alignment with spatial RNA data. During permeabilization, mRNA was liberated from cells and captured by primers on the slide’s surface which enable downstream reassignment of barcoded mRNA sequences to their former, spatial location. Permeabilization times had been determined in advance (Cortex: 18 min; Tumor: 12 min; Cerebellum: 12 min) according to manufacturer’s instructions (10X Genomics, Spatial Tissue Optimization Reagent Kit). After reverse transcription, second strand synthesis and denaturation of cDNA, second strands were amplified by PCR and desired cDNA fragments were selected via SPRIselect reagent. Successful amplification was confirmed by QC via Agilent Fragment Analyzer system. During the following fragmentation and double-sided size selection via SPRIselect reagent, length of cDNA fragments was optimized for analysis via Illumina NextSeq Sequencing System. Each fragment was provided with unique, dual indexes as well as adapters binding to oligonucleotides on Ilumina flow cell. Post Library Construction QC via Agilent Fragment Analyzer system and Invitrogen Qubit Fluorometer was performed before normalization of libraries. For more information consult Illumina ‘Denature and Dilute Libraries Guide - Protocol A: Standard Normalization Method’. Phix control at a concentration of 1.8pM was added to each library in a dilution of 1:100. Sequencing was performed using the NextSeq 500/550 High Output Kit (150 Cycles).

### Data and code Availability

Further information and requests for resources, raw data and reagents should be directed and will be fulfilled by the Contact: D. H. Heiland. The source code of SPATA is available at https://github.com/theMILOlab/SPATA, additional functions are at https://github.com/heilandd/SPATA_Developer and https://github.com/kueckelj/confuns. Spatial Transcriptomic data will be provided at GEO (in preparation) and SPATAobjects at www.themilolab.com (in preperation).

## Acknowledgement

This study is funded by the Else Kröner-Fresenius-Stiftung (2020_EKSMS.24) (DHH), BMBF (MEPHISTO) (DHH and DD) and the Krebshilfe (Mildred-Scheel-Program) (JK and JE)

